# Inverse reinforcement learning reveals action-oriented value signals in naturalistic decision making

**DOI:** 10.64898/2026.06.24.733779

**Authors:** Sang Ho Lee, Chaeyoun Chung, Min-hwan Oh, Woo-Young Ahn

## Abstract

A major challenge for cognitive neuroscience is to explain how value of a goal-directed behavior is computed in complex and naturalistic environments. Standard computational models of decision making have been highly successful in controlled, trial-based paradigms, but they are often ill-suited to real-time behavior unfolding in naturalistic paradigms. Inverse reinforcement learning (IRL) offers a way to infer latent evaluative state from observed behavior in naturalistic environments, but its neural interpretability remains largely unknown. Here, we investigated whether moment-to-moment reward trajectories derived from IRL map onto value signals in the brain during a real-time driving task performed during fMRI scanning. IRL-derived reward trajectories were most robustly associated with activity in the dorsal striatum, a region often linked to value-guided action selection. They also showed associations with distributed regions supporting additional processes, including cognitive control and sensorimotor processing. This pattern suggests that IRL reward captures distributed neural activity centered on the reward circuitry, potentially reflecting how valuation interacts with other processes. Together, these findings suggest that IRL reward provides a behaviorally grounded, temporally resolved proxy for action-oriented valuation during naturalistic decision making.

## Introduction

Understanding the mechanisms underlying real-life decision making has long been a central question in cognitive neuroscience. Historically, the field has relied on highly controlled, trial-based laboratory tasks designed to isolate specific cognitive processes^1–4^. While these paradigms have yielded foundational insights, the simplicity of such paradigms raises concerns about whether the insights derived from the tasks extend to the complexities of real-world decision making^5–7^. Recent studies address this limitation by using naturalistic tasks that better capture the dynamics of everyday behavior, such as those implemented in virtual reality^8–10^ and video games^11,12^. These paradigms improve ecological validity by immersing participants in complex, realistic environments where they make decisions in real time.

A central component of real-life decision making is evaluating the value of available actions given the current state^13,14^. For example, when driving in traffic, a driver must continuously decide whether to speed or brake based on the movements of surrounding vehicles. The relative values of speeding and braking, which continuously change with the current state, determine which action is more likely to be selected. Understanding how such value estimates fluctuate in real-world settings is therefore critical for explaining naturalistic behavior.

However, investigating these dynamics is inherently challenging because internal value is not directly observable. This limitation has motivated the use of computational models to infer latent subjective values from observed behavior^2,15^. Developing such models typically requires the researcher to manually define a reward function over all relevant states and events in the task. This is feasible in highly controlled, trial-based tasks, but particularly challenging in real-time tasks with countless states, which are defined by a combination of multiple variables such as spatial orientation and velocity of many surrounding objects.

Alternatively, highly flexible data-driven models, such as deep Q-networks (DQN)^16^, can effectively model the behaviors in high-dimensional environments by learning direct mappings from complex sensory inputs to motor outputs^17^. However, these models often operate as black boxes, offering limited interpretability and little insight into the latent cognitive variables that guide decision making and explain individual differences. For example, DQN are trained to maximize task performance rather than to model human behavior, making them ill-suited for explaining individual differences. This creates a fundamental challenge in naturalistic decision-making research: conventional computational models are interpretable but difficult to scale to complex real-time environments, whereas purely data-driven models can handle such complexity but sacrifice explanatory power.

Inverse reinforcement learning (IRL)^18,19^ provides a compelling synthesis of interpretable computational modeling and data-driven techniques. While conventional forward reinforcement learning seeks to determine the optimal action strategy (policy) given a predefined reward function, IRL infers the underlying reward function by observing behaviors. By using a trajectory of state and action as the input, IRL can approximate latent reward for every state and action within a high-dimensional space. The IRL-derived reward then serves as a computational proxy to be linked with cognitive traits or corresponding brain signals. For example, Lee et al. showed that IRL can successfully model human behavior in a real-time driving task and recover the reward trajectories associated with impulsivity^20^.

A central assumption underlying the use of IRL to model behavioral data is that the inferred reward reflects how favorable the current state is for ongoing action. This raises an important question: does IRL-derived reward correspond to value signals in the brain? If moment-to-moment fluctuations in IRL reward track changes in BOLD activity within the reward-related circuitry such as the striatum^21^, this would support its interpretation as a proxy for internal value.

At the same time, because IRL reward is inferred from behavior, it may also relate to activity in other systems that support continuous action selection. For example, sensorimotor regions (e.g., supplementary motor area) may reflect the execution of action^22^, which is directly linked to the behavioral data used for inference. Control-related regions such as the anterior cingulate cortex (ACC) may also track the need to adjust behavior to optimize ongoing performance^23^.

The present study addresses this question in an fMRI experiment using a real-time driving task in which participants control a car on a simulated highway as quickly as possible while avoiding collisions^20^. We assess the neurobiological plausibility of the IRL framework by examining whether the IRL reward trajectories track moment-to-moment brain activity, with a particular focus on the reward-related circuitry and additional analyses of cognitive control and sensorimotor systems^24^.

## Results

### Neural responses in the reward circuit during the highway task

Prior to testing the association between IRL reward and BOLD signals, we evaluated whether the highway task engaged the reward circuit. This step established that the task elicited meaningful value signals, thereby providing an appropriate foundation for evaluating the neural interpretability of IRL in the subsequent analyses.

General linear model (GLM) analysis examined BOLD signals associated with two salient task events, overtaking and crash (**Fig. 1b-c**), using event-onset regressors. The striatum, which is a key region in the reward circuit^25–28^, showed robust task-related responses. **Fig. 2** shows whole-brain t-statistic maps displayed at the peak coordinates of clusters that include the striatum. Overtaking was associated with increased activation in the striatum, whereas crash was associated with decreased activation in the striatum. In addition to the reward circuit, activation was observed across occipital, parietal, cerebellar, and sensorimotor cortices (see **Supplementary Table 5** for the full list of clusters and statistics), consistent with the visuomotor demands of the task.

**Fig. 1.**
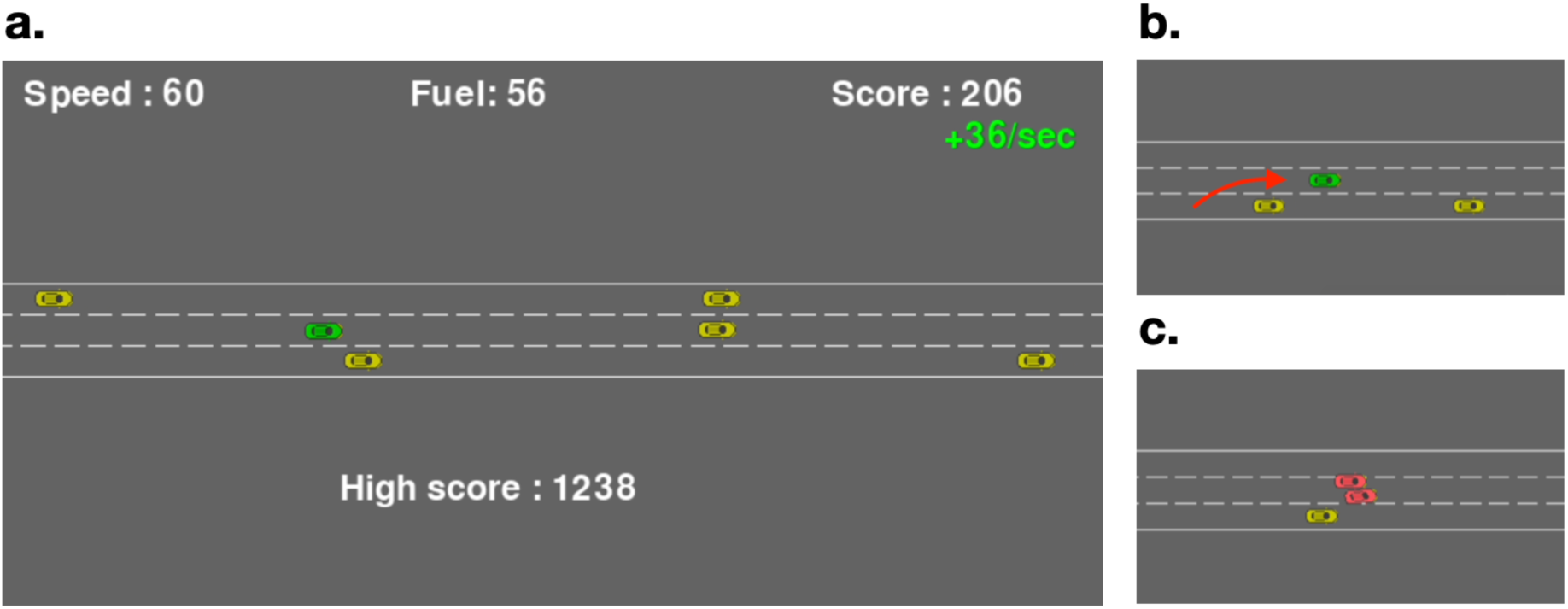
| **The highway task.** (a) Screenshot of the task interface. (b, c) Two salient events in the task: b) *overtaking,* in which the participant’s vehicle passes another vehicle, and c) *crash,* in which the participant’s vehicle collides with another vehicle.

**Fig. 2.**
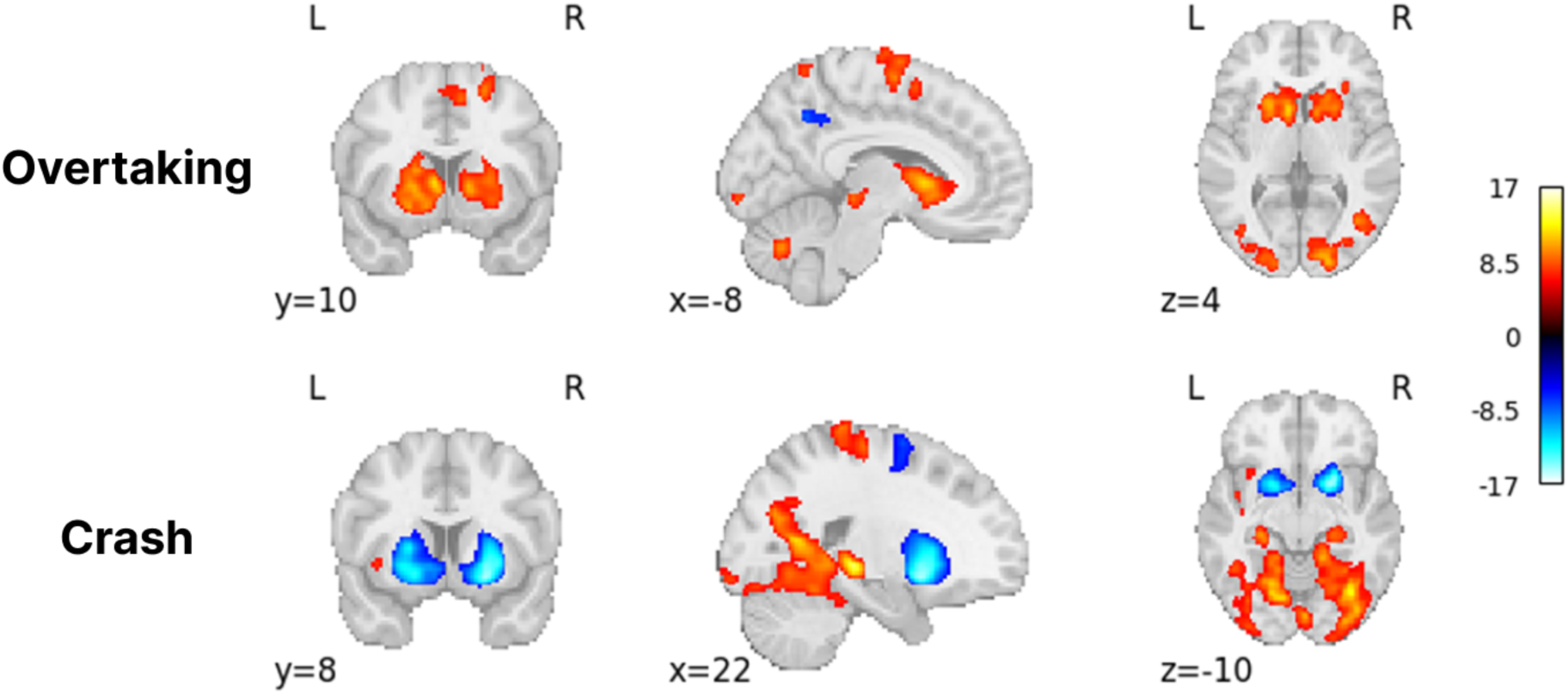
| Whole-brain t-statistic maps (Bonferroni-corrected *p*<0.01) for the neural responses to the onsets of overtaking or crash events.

### IRL model evaluation and behavioral data analysis

Having established that the highway task engaged robust neural responses in the striatum, we next evaluated the performance of the IRL model, as a necessary step before testing the neural interpretability of IRL reward. The behavioral policy learned by the algorithm predicted the observed actions with high accuracy (total accuracy = 0.86, above chance level for all actions; **Fig. 3a**; see Methods for details), with predicted action frequencies closely resembling those observed in the data (**Fig. 3b**), indicating that the IRL model provided a good fit to the behavioral data. The IRL reward map was broadly consistent with that reported in Lee et al. (2024)^20^. IRL reward generally decreased with speed and increased with distance from the car ahead (**Fig. 3d**). More specifically, reward tended to be high at low speed and moderate distance, and low at high speed and short distance (**Fig.3c**), reflecting a generally safe driving strategy. This safety-oriented strategy was further evidenced by the rarity of high-speed states (>100), which comprised only 0.4% of the total dataset (See **Supplementary Fig. 2** for details). These states were omitted from the visualization (**Fig. 3c-d**), as the scarcity of observations in those regions of the state space precluded stable reward estimation.

**Fig. 3.**
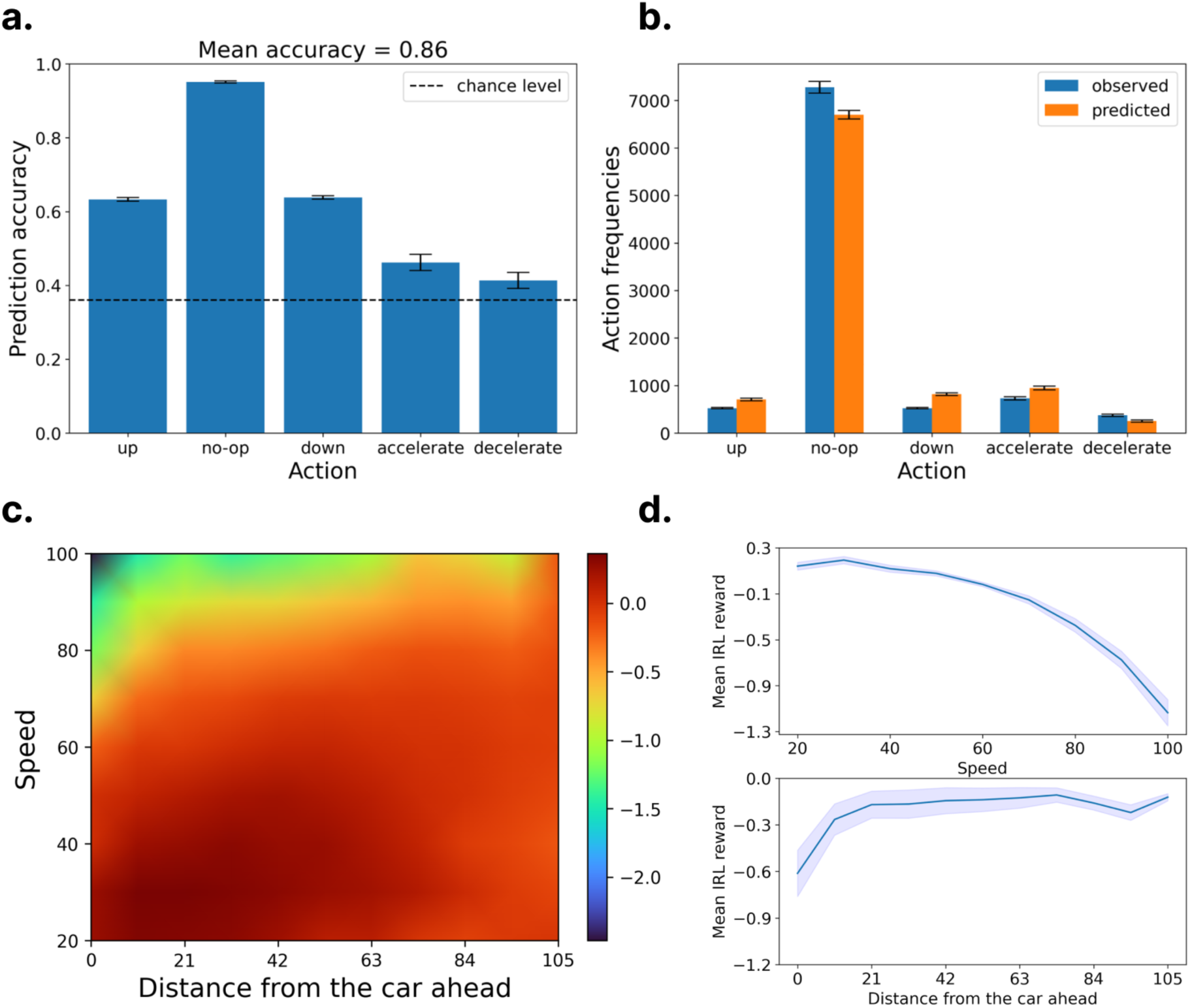
| **IRL model evaluation and recovered reward structure. (a)** Action prediction accuracy of the model. **(b)** Histogram comparing observed and predicted action frequencies. **(c)** Heatmap of the mean IRL reward as a function of speed and distance from the car ahead. **(d)** Mean IRL reward functions across speed and distance dimensions.

To assess whether task performance was influenced by individual differences beyond general task strategy, we examined its association with participant background and traits (see **Supplementary Table 1-4, Supplementary Fig. 1** for details). Among the factors assessed, including motor skills, impulsivity, and prior experience, none showed a significant correlation with task score. This suggests that the IRL framework may capture task-specific valuation independent of general motor or experiential confounds, which are difficult to disentangle in real-time tasks^29^.

### Temporal alignment between IRL-derived reward and BOLD signals

#### Full-trajectory analysis

The primary hypothesis of this study is that the reward inferred by IRL reflects endogenous reward processing in the brain. To test this hypothesis, we first evaluated whether BOLD signals across the whole brain could reconstruct the IRL reward trajectory throughout the data. The IRL reward was computed at a high temporal resolution (0.2 s) and aligned to the fMRI sampling rate (TR = 1.2 s), yielding a total of 66,224 time points across all participants (n = 45).

A linear mixed-effects model with Elastic Net penalty (see Methods for model specification) was employed to predict the IRL reward from BOLD signals across 120 ROIs in AAL2 atlas^30^ while controlling for subject-level random effects. Under a leave-one-participant-out cross-validation scheme, the model demonstrated a robust correlation between the predicted and observed IRL reward trajectories (*r* = 0.37; *p* < .001). Given the size and complexity of the dataset, this stable association indicates that the IRL reward consistently reflects neural fluctuations throughout the task.

The full-trajectory analysis revealed a distributed pattern of contributions across ROIs, with Elastic Net retaining a broad set of regions. **Fig. 4a** shows the whole-brain map of beta coefficients for the ROIs retained by Elastic Net, illustrating the relative contribution of each region to the reconstruction of the IRL reward. Regions with the highest contributions (top 10% of absolute beta coefficients) included middle frontal gyrus (MFG), precentral gyrus, putamen, visual cortex (fusiform, calcarine, occipital), hippocampus, and cerebellum (see **Supplementary Table 6** for the full list of beta coefficients).

**Fig. 4.**
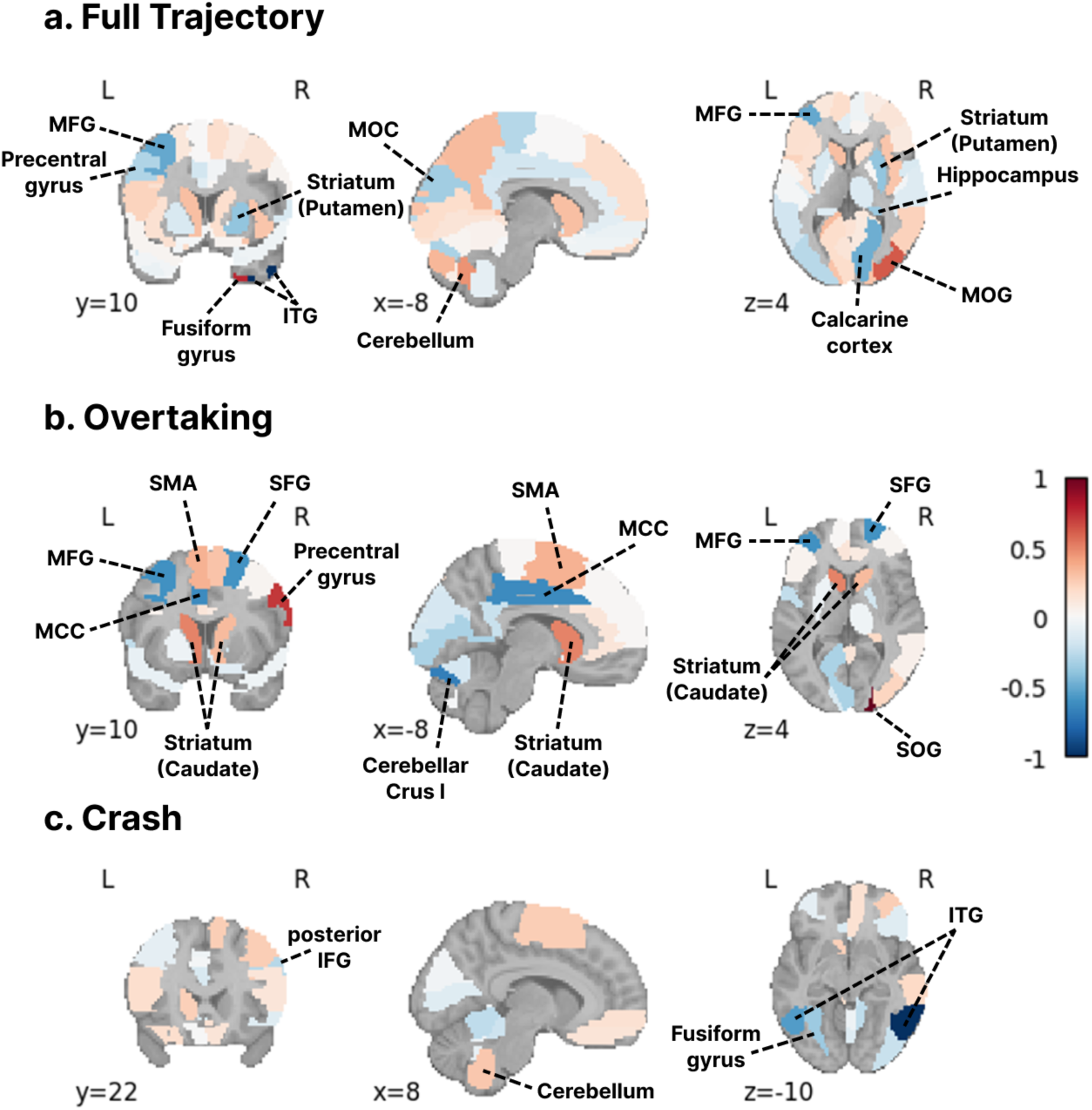
| Beta coefficient maps for neural prediction of IRL reward. ROIs defined by the AAL2 atlas are color-coded to indicate their contributions. Anatomical distributions of beta coefficients are shown for the models predicting a) full IRL reward trajectory, b) IRL reward within a ±10-TR (12 s) window centered on overtaking events, and c) IRL reward within a ±10-TR window centered on crash events. ROIs meeting threshold criteria (top 10% for full-trajectory and top 25% for event-based analyses) are labeled with anatomical names. Abbreviations: MFG, middle frontal gyrus; MOC, medial occipital cortex; ITG, inferior temporal gyrus; MOG, middle occipital gyrus; SMA, supplementary motor area; SFG, superior frontal gyrus; MCC, mid-cingulate cortex; IFG, inferior frontal gyrus.

#### Event-level analysis

The highly distributed pattern observed in the full-trajectory analysis motivated a more temporally focused approach. In light of GLM results indicating clearer event-related contrasts, particularly in the striatum, we next tested whether an event-based analysis would better specify the ROIs associated with IRL reward.

The mapping between BOLD signals and the IRL reward was markedly more precise when the analysis was restricted to specific events in the task (i.e. overtaking and crash). In the event-level analysis, the model predicted the IRL reward trajectory within a ±10-TR window (12 s) centered on event onset. The BOLD signals and the IRL reward trajectories were averaged across events within each participant to obtain participant-level event time series (see Methods for the details). The IRL reward predicted by the model with ROIs defined by AAL2 atlas showed a strong correlation with the observed reward trajectory (overtaking *r* = 0.86, *p* < .001; crash *r* = 0.81, *p* < .001). Parallel analyses using only the reward-specific ROI defined via Neurosynth mirrored these results (overtaking *r* = 0.80, *p* < .001; crash *r* = 0.76, *p* < .001), suggesting that the model’s predictive power is consistently anchored in reward-related regions.

The anatomical distributions of beta coefficients are illustrated in **Fig. 4b** and **4c** for overtaking and crash events, respectively. ROIs within the top 25% of absolute beta coefficients are labeled in the figures; a more inclusive threshold (25%) was used relative to the full-trajectory analysis (10%) because fewer ROIs were retained by Elastic Net. For overtaking events, strong contributions were distributed across frontal, striatal, occipital, sensorimotor, and cerebellar regions. This pattern is consistent with the GLM results, which showed robust activation in the striatum alongside occipital, parietal, sensorimotor, and cerebellar cortices. For crash events, fewer ROIs showed robust associations with IRL reward, with contributions primarily concentrated in occipital, temporal, parietal, and cerebellar regions.

#### IRL reward and BOLD trajectory by ROI

The event-level analysis identified ROIs exhibiting robust associations with the IRL reward. For ROIs that survived the variable selection of the Elastic Net, changes in the BOLD signals were compared with the IRL reward trajectories, with Pearson’s *r* quantifying the similarity between the two trajectories. For this comparison, IRL reward trajectories were convolved with a hemodynamic response function. The BOLD signal and IRL reward trajectories were averaged across subjects and z-normalized. BOLD signals from many of the selected regions closely tracked the IRL reward trajectory, showing robust correlations (Bonferroni-corrected *p* < 0.01) in 28 ROIs for overtaking and 12 ROIs for crash (**Table 1**).

**Table 1:** Correlations between IRL reward and BOLD signals in selected ROIs.

| ROI | Overtaking (left/right $r$) | Crash (left/right $r$) |
| --- | --- | --- |
| Middle frontal gyrus | left: -0.82 | — |
| Supplementary motor area | 0.83 / 0.88 | — |
| Medial superior frontal gyrus | left: -0.90 | left: -0.83 |
| Anterior cingulate cortex | -0.90 / -0.82 | left: -0.93 |
| Posterior cingulate cortex | right: -0.77 | — |
| Hippocampus | right: -0.87 | — |
| Calcarine cortex | left: -0.86 | — |
| Cuneus | left: -0.77 | -0.94 / -0.86 |
| Superior occipital gyrus | 0.96 / 0.90 | — |
| Middle occipital gyrus | right: 0.88 | — |
| Inferior occipital gyrus | — | right: -0.84 |
| Superior parietal lobule | 0.89 / 0.92 | left: 0.93 |
| Angular gyrus | right: -0.93 | — |
| Caudate | 0.84 / 0.83 | left: 0.90 |
| Thalamus | right: 0.86 | — |
| Superior temporal pole | right: -0.90 | — |
| Inferior temporal gyrus | — | -0.89 / -0.87 |
| Rolandic operculum | — | right: -0.83 |
| Cerebellar Crus I | left: -0.80 | — |
| Cerebellum lobules VI, VIIb | left: 0.93, 0.86 | — |
| Cerebellum lobules IX, X | right: 0.83, 0.93 | right: 0.94, 0.89 |
| Vermal lobules IV–V, VI, VII | 0.94, 0.88, 0.82 | — |

Beyond the striatum (caudate), correlations were observed in a broad set of regions, including those associated with cognitive control (e.g., middle frontal gyrus (MFG)^31^, anterior cingulate cortex (ACC)^23^ and sensorimotor processing (e.g., supplementary motor area (SMA)^32^). Notably, control-related regions showed predominantly negative correlations, whereas sensorimotor regions showed strong positive correlations.

To further support these findings, we conducted analyses using meta-analytically defined functional ROIs from Neurosynth (https://neurosynth.org). The ROIs were derived from association maps for the terms “reward” (922 studies), “cognitive control” (598 studies), and “sensorimotor” (684 studies). **Fig. 5** shows the resulting ROI maps and BOLD time series from these ROIs overlaid on the HRF-convolved IRL reward trajectories during overtaking and crash events. The reward ROI showed moderate positive correlations (overtaking r = 0.43; crash r = 0.66). The cognitive-control ROI showed weak correlations (overtaking r = 0.03; crash r = −0.15), potentially because of the relatively sparse and spatially restricted nature of the meta-analytic map (**Fig. 5b**). In contrast, the sensorimotor ROI exhibited strong positive correlations (overtaking r = 0.87; crash r = 0.64).

**Fig. 5.**
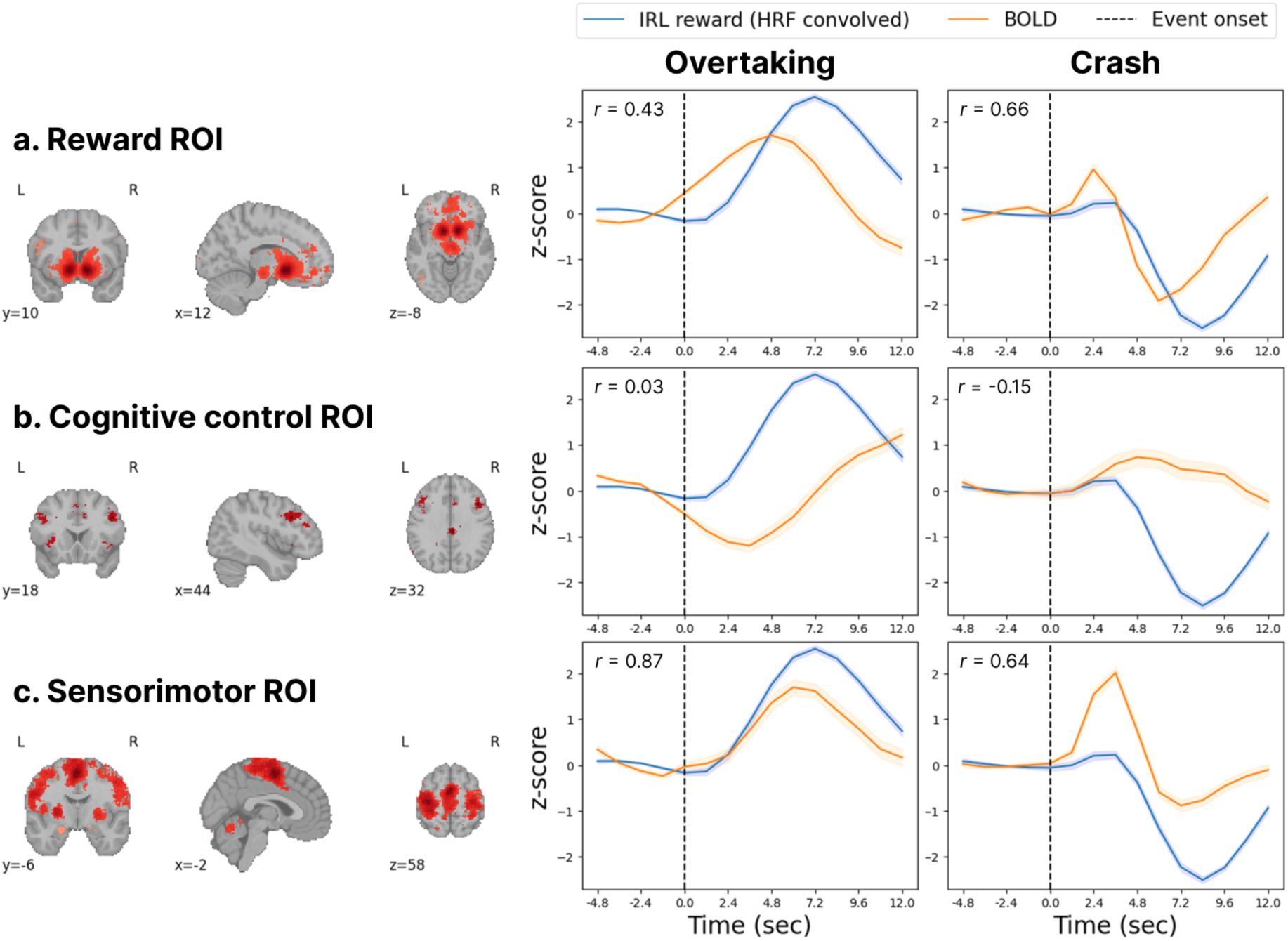
| Correlation analysis using meta-analytically defined ROIs from Neurosynth. Left panels illustrate ROI maps for the terms (a) reward, (b) cognitive control, and (c) sensorimotor. Right panels show BOLD time series from these ROIs overlaid with IRL reward trajectories. Time series were baseline-corrected by subtracting the mean signal from −4 TR (4.8 s) to event onset, setting the pre-event baseline to zero.

Correlations were much stronger in specific ROIs defined by the AAL2 atlas, where BOLD signals closely tracked changes in IRL reward. **Fig. 6** illustrates BOLD time series from representative ROIs with particularly strong correlations with IRL reward. Strong positive correlations were observed in the striatum (caudate; overtaking r = 0.84, crash r = 0.90), whereas the ACC showed strong negative correlations (overtaking r = -0.90, crash r = -0.93). The SMA showed strong positive correlations, with a lower correlation during crash events (overtaking r = 0.88; crash r = 0.71).

**Fig. 6.**
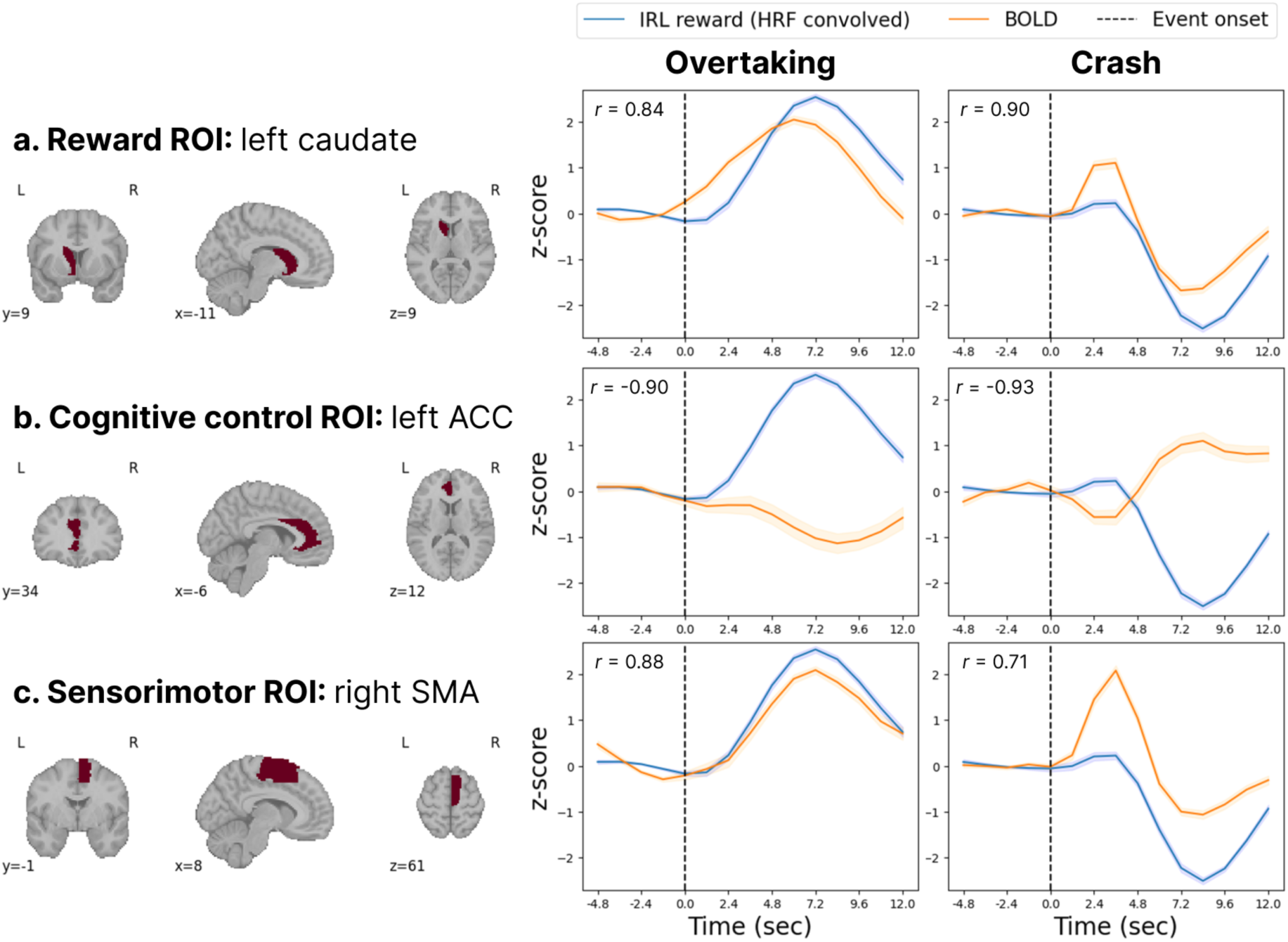
| Correlation analysis using selected AAL2 ROIs. BOLD time series from selected AAL2 ROIs were overlaid with HRF-convolved IRL reward trajectories. Time series were baseline-corrected relative to the pre-event period, as in Fig. 5.

As an additional specificity analysis, we tested whether the IRL reward could be replaced by the Neurosynth-based *reward* BOLD trajectory. When this BOLD trajectory was used in place of IRL reward for correlation analyses with other regions, the number of ROIs showing robust correlations was markedly reduced (see **Supplementary Table 7** for the full list of correlation coefficients). This suggests that the regional correlation pattern obtained with IRL reward is distinct from that obtained using a BOLD trajectory derived from the reward circuit alone.

## Discussion

The present study demonstrates the utility of IRL reward as a behaviorally grounded computational proxy for neural dynamics associated with ongoing task-state evaluation. Across analyses, the IRL reward showed meaningful correspondence with BOLD activity, from whole-trajectory prediction across the task to event-level prediction around overtaking and crash events. These findings suggest that IRL can provide a temporally resolved estimate of latent evaluative state that is neurally interpretable, even in a complex, naturalistic task.

The highway task successfully elicited neural responses in the reward circuit. Overtaking events, which were associated with rewarding outcomes, produced increased activation within the striatum, whereas crash events, corresponding to punishing outcomes, were associated with reduced striatal activation. This dissociation aligns with the reward structure of the task and validates the paradigm’s capacity to capture value signals in a naturalistic setting.

### Neural Correlates of IRL-derived Reward

The time-series analysis showed that distributed BOLD activity across 120 ROIs reliably tracked fluctuations in IRL reward. Engagement of the striatum was remarkably consistent across analyses. During overtaking, the bilateral caudate showed robust positive correlations with IRL reward (left: r = 0.84; right: r = 0.83), characterized by a striking temporal coupling between the IRL reward trajectory and the BOLD time series (**Fig. 6**). A similar positive association was found in the left caudate during crashes (r = 0.90). This pattern is consistent with the GLM results, which showed a clear contrast in striatal activity between overtaking and crash events. The finding that caudate activity tracked IRL reward in both successful and adverse task contexts suggests that the IRL reward consistently captured fluctuations in neural valuation signals, rather than merely reflecting rewarding outcomes.

A more precise interpretation of these findings can be informed by the functional distinction between dorsal and ventral striatum. Caudate largely corresponds to the dorsal striatum^33^, which has been implicated in goal-directed action selection and action-outcome associations^27^. On the other hand, the ventral striatum has been more commonly associated with reward anticipation^34^. In this framework, the predominance of caudate involvement in the present results suggests that IRL-derived reward reflects action-contingent valuation rather than purely Pavlovian reward anticipation or cue-evoked value signals.

#### Cognitive Control and Sensorimotor Processing

The correlations with IRL reward were not limited to the striatum but extended to distributed regions. A notable finding is the strong inverse relationship between IRL reward and cognitive control regions. For overtaking events, regions associated with cognitive control, including bilateral ACC^23^, left MFG^31^, and left cerebellar Crus I^35^ exhibited robust negative correlations with IRL reward. A similar pattern was observed for crash events, with a negative correlation in the left ACC (*r* = -0.93).

The association between ACC and IRL reward suggests that the brain may dynamically regulate deliberative, effortful cognitive control as a function of task-state valuation. This interpretation aligns with the expected value of control model^36^, which posits that the ACC dynamically regulates executive investment based on a cost-benefit analysis of the current task state. In this context, higher IRL reward may index states in which the agent is effectively meeting its behavioral objectives, reducing the utility of sustained deliberative control. Notably, this region-specific pattern contrasts with the weak correlations observed for the meta-analytic cognitive-control ROI, likely reflecting differences in spatial specificity: the broadly distributed Neurosynth ROI may dilute region-specific effects, whereas focal regions such as the ACC capture more specific associations with IRL reward.

In contrast, sensorimotor regions, such as the bilateral SMA (r = 0.83/0.88), showed strong positive correlations with IRL reward. This pattern suggests that action execution during the task is closely aligned with higher-value states, such that ongoing behavior naturally unfolds in directions that maximize reward.

Activity in these regions may reflect the implementation of behavior shaped by the underlying value of actions, rather than the valuation process itself.

Taken together, these findings suggest that IRL reward captures a distributed neural representation of task-state value that extends beyond the reward circuit. In this framework, IRL reward serves as a temporally resolved computational proxy for ongoing, action-oriented valuation, reflecting how favorable the current state is for goal-directed behavior.

#### Comparison of IRL reward and reward-circuit BOLD signals

When IRL reward was replaced with a Neurosynth-based reward BOLD trajectory in the correlation analysis, correlations across ROIs were altered and attenuated (**Supplementary Table 7**), suggesting that IRL reward is not simply interchangeable with the BOLD signals from the reward circuit alone. Crucially, IRL reward is inferred from behavioral trajectories and thus provides a behaviorally grounded computational proxy for task-state valuation. Replacing IRL reward with a BOLD trajectory changes the analysis from testing how behaviorally inferred valuation relates to neural activity to testing how one neural time series covaries with another. The distinct association between IRL reward and BOLD activity, relative to BOLD-to-BOLD correlations, suggests that IRL reward may reflect action-oriented valuation in a way that complements conventional connectivity analyses.

### Limitation and future research

While the results of the current study provide important insights, two limitations should be considered. First, the observed relationships between IRL reward and BOLD signals are correlational. Future work is needed to establish the causal role of specific cognitive processes in modulating valuation signals captured by IRL. For example, experimental manipulations that vary task difficulty or uncertainty may modulate control demands and help clarify how IRL reward relates to the engagement of cognitive control processes. Second, while IRL provides a useful framework for modeling behavior in complex tasks, performance in naturalistic settings may also be strongly influenced by motor control demands. For example, increased control difficulty in the fMRI environment may encourage more conservative or safer behavioral strategies, which could in turn influence the reward function inferred by IRL. This tendency is reflected in the current data, where high-speed states (>100) were extremely rare.

### Conclusion

The present study demonstrates that IRL reward provides a behaviorally grounded and temporally resolved proxy for neural valuation signals in a complex, naturalistic task. By linking behavioral trajectories to brain activity, we show that IRL reward corresponds with activity in distributed regions associated with task-state valuation. These findings highlight the potential of IRL as a framework for bridging behavior and neural representations of value in naturalistic decision making.

## Methods

### Participants

A total of 50 undergraduate and graduate students were initially recruited to participate in the experiment at Seoul National University fMRI Center. Five participants were excluded from analysis due to the following reasons: fMRI scanner malfunction (n=2), poor data quality due to drowsiness during the task (n=1), and preprocessing failure due to incorrectly saved imaging files (n=2). The final sample for the primary analysis thus comprised 45 participants (22 males, 23 females; age = 20.22 ± 2.84 years). One additional participant was excluded from finger tapping analysis, and another participant was excluded from gaming survey due to missing data.

### Experiment

Participants first completed a Korean version of the Barratt Impulsiveness Scale (BIS)^37,38^, a 30-item self-report questionnaire designed to assess multiple dimensions of impulsivity. An experimenter then provided instructions for the highway task^20^, followed by a 5-minute practice session. Participants subsequently performed the highway task during fMRI scanning for approximately 40 minutes, divided into four 10-minute blocks. They were given the opportunity to take a break after each block for as long as they desired.

Following the highway task, participants completed a finger-tapping task and a post-experimental survey outside the scanner. Finger-tapping task performance assessed motor control ability, whereas survey responses assessed prior gaming and driving experience as potential confounds of highway task performance.

#### Highway task

Participants were instructed to drive a car on a simulated highway as fast as possible without crashing into other cars. The car was controlled via a four-button response box, with each button corresponding to one of four possible actions: accelerate, decelerate, turn left, and turn right. For a detailed explanation of the task, see ref.^20^

#### Finger-tapping task

To assess baseline motor proficiency, participants performed a sequential finger-tapping task^39^ outside the fMRI scanner. Using the non-dominant hand, participants positioned their fingers (index to little) on the numerical keys (1–4) of a keyboard. The task required the repeated execution of a five-element sequence (4-1-3-2-4) as rapidly and accurately as possible within 30-second blocks. The experiment consisted of 12 trials, each interspersed with 30-second rest periods to minimize fatigue.

#### Post-experimental survey

The survey assessed participants’ driving experience and gaming experience. Driving experience was measured along two dimensions: frequency and duration. Driving frequency was assessed using a 5-point ordinal scale: *less than once per month, less than once in a week - more than once in a month, 1-2 days per week, 3-4 days per week, 5 or more days per week.* Driving duration, defined as the total length of driving experience, was rated on a 6-point ordinal scale: *less than 6 months, 6 months to one year, 1-3 years, 3-5 years, 5-10 years, more than 10 years.* Gaming experience was assessed across five genres, including racing, sports, action, role-playing, and other, along two dimensions. Gaming frequency was measured on a 3-point ordinal scale: *every day, every week, less than once per month.* Participants also reported the typical duration of a single gaming session on a 5-point ordinal scale: *0-1 hour, 1-2 hours, 3-5 hours, 5-10 hours, more than 10 hours*.

### Behavioral analysis

#### Task event specification

The primary events of interest in the highway task were overtaking and crash. Overtaking was defined as the moment the participant’s vehicle passed computer-controlled cars. Crash was defined as a collision between the participant’s vehicle and computer-controlled cars.

#### Inverse Reinforcement Learning (IRL)

We employed Adversarial Inverse Reinforcement Learning (AIRL)^40^, which utilizes a generative adversarial network (GAN)^41^ framework to infer reward functions from observed behaviors. AIRL models learning as an adversarial process between a generator, which learns a policy that reproduces expert-like behavior, and a discriminator, which distinguishes expert behavior from generated behavior. Both components were implemented as deep neural networks to capture nonlinear relationships among states, actions, and rewards, enabling the recovery of complex, high-dimensional reward functions. For details of the model specification, see the codes and the instructions on the GitHub repository, which will be publicly available upon the publication of the paper.

#### Policy accuracy calculation

To calculate the accuracy of the policy learned through AIRL (**Fig. 2a**), we computed a lag-adjusted accuracy measure. A predicted action was counted as correct if it matched the observed action at the same time point or one time step earlier, to allow for delayed behavioral execution relative to the model-predicted action. This adjustment was motivated by the possibility that overt motor actions may occur slightly later than the underlying intention, particularly in the MRI environment where the constrained posture and use of an MRI-compatible button box may make responses less comfortable and less temporally precise than those made on a keyboard. As a reference, we used a random baseline assuming uniform guessing across five actions. Because a prediction was counted as correct if it matched at the current or subsequent time step, the corresponding random baseline was 1 − (1 − 0.2)^!^ = 0.36, reflecting the probability of at least one match across two time points.

### Neuroimaging data acquisition and analysis

#### Data acquisition

All MRI data were acquired at the Seoul National University Brain Imaging Center using a Siemens TIM Trio 3T scanner (Siemens Healthineers, Erlangen, Germany) equipped with a 32-channel head coil. Functional images were collected using a T2*-weighted multiband gradient-echo EPI sequence with the following parameters: TR = 1200 ms, TE = 30 ms, flip angle = 85°, voxel size = 2.3 x 2.3 x 2.3 mm^3^, slice thickness = 2.3mm, FOV = 256mm, matrix size = 110×110, 64 axial slices, multiband acceleration factor = 4. In addition, a high-resolution T1-weighted structural image was acquired using a 3D MPRAGE sequence with the following parameters: TR = 2300 ms, TE = 2.36 ms, TI = 700 ms, flip angle = 9°, voxel size = 1.0 x 1.0 x 1.0 mm^3^, 224 sagittal slices.

#### Data preprocessing

fMRI data were preprocessed using fMRIPrep (version 20.2.6)^42^, which is based on Nipype 1.7.0. Preprocessing included motion correction, co-registration to the T1-weighted reference, and normalization to MNI152NLin2009cAsym standard space. Functional images were spatially smoothed with a 6-mm FWHM Gaussian kernel prior to the analysis.

#### Data analysis

Statistical analyses were conducted using in-house Python code with the nilearn and nltools packages (GitHub repository will be publicly available upon the publication of the paper).

*General linear model (GLM) analysis.* First-level voxel-wise whole-brain GLMs were estimated for each participant using ordinary least squares. The design matrix included five task regressors reflecting driving behaviors (lane changing, acceleration, deceleration, crash and overtaking), each convolved with a hemodynamic response function, and six head-motion nuisance regressors, comprising three rotational and three translational parameters.

In the second-level analysis, voxel-wise one-sample t-tests were performed on the first-level contrast. The main contrasts of interest were overtaking onset and crash onset. The resulting voxel-wise *t*-statistic maps were corrected for multiple comparisons using a Bonferroni correction (corrected *p* < 0.01). Clusters exceeding this threshold were identified.

*Time-series analysis.* We examined the temporal correspondence between the reward estimates derived from IRL and BOLD signals throughout the task. To account for the hemodynamic delay of the BOLD response, the IRL-derived reward was convolved with a hemodynamic response function using the nltools package in Python. The BOLD time series for each region of interest (ROI) was defined as the mean BOLD signal across all voxels within the region. ROIs were defined by AAL2 atlas^30^ and Neurosynth parcellations^43^.

We combined a linear mixed-effects model with regularized regression (Elastic Net) to predict IRL-derived reward trajectories while accounting for between-subject variability. We first fit a linear mixed-effects model with subject-specific random intercepts and removed these effects from the IRL reward trajectory. The residualized signal was then predicted using Elastic Net regression with leave-one-participant-out cross-validation, where data from each participant were held out in turn. This approach captures time-varying neural contributions while preventing participant-level leakage and overfitting.

Hyperparameters of the Elastic Net, including the regularization strength (α) and the L1–L2 mixing parameter (l1_ratio), were selected via cross-validation by evaluating multiple candidate values and choosing those that minimized prediction error. Potential confounding variables, including the timing of action execution (i.e., acceleration, deceleration, and lane changing), were included in the model as covariates. Participant-specific random intercepts were included to account for inter-individual differences in baseline IRL reward.

In the full-trajectory analysis, the entire time series from all participants was concatenated to form a continuous trajectory. In the event-based analysis, task-relevant events (e.g., overtaking and crash events) were extracted, and BOLD signals or IRL reward estimates were averaged within event-specific time windows (±10TR).

## Data and Code availability

The data generated in this study will be made available in an Open Science Framework repository upon publication. The analysis code will be made available in a GitHub repository. Additional information required to reproduce or interpret the findings is available from the corresponding author upon request.

## Author Contributions

S.H.L., M.O., and W-Y.A. conceived and designed the research. S.H.L. and C.C. collected and analyzed the data. M.O. supervised the methodological development. W-Y.A. supervised the interpretation of the results. S.H.L. and C.C. wrote the initial draft. All authors reviewed and edited the manuscript and approved the final version.

