## Supplementary Information for "Inverse reinforcement learning reveals action-oriented value signals in naturalistic decision making"

#### Supplementary Results

##### Association between prior experience and task performance

We conducted exploratory analysis of the post-experimental survey to examine associations between prior driving or gaming experience and highway task performance. Data from a total 44 participants were included in the analysis.

Survey responses were systematically coded. Binary items were coded as 1 (*yes*) and 0 (*no*), while frequency and duration items were coded on ordinal numeric scales reflecting increasing levels (e.g., driving frequency: 1 = *less than once a month*, 5 = *nearly every day*; gaming frequency: 1 = *less than once per month*, 3 = *every day*). Open-ended responses were manually assigned to the appropriate category. Group differences based on binary experience were assessed using Welch's t-tests. We found no significant difference in task scores between participants with ($n$ = 21, $M$ = 872.93, $SD$ = 222.98) and without ($n$ = 23, $M$ = 845.71, $SD$ = 199.68) driving experience, $t$(40.35) = 0.425, $p$ = 0.673. Similarly, scores did not significantly differ between participants with ($n$ = 34, $M$ = 873.37, $SD$ = 217.28) and without ($n$ = 10, $M$ = 808.83, $SD$ = 155.38) gaming experience, $t$(42) = 1.01, $p$ = 0.33. Further comparisons by specific gaming genres revealed no significant differences in task performance between participants with and without experience in a given genre (with experience: Action: $n$ = 34, Racing: $n$ = 32, Sports: $n$ = 32, Role-playing: $n$ = 36, Others: $n$ = 19; all $p$s > 0.34; **Supplementary Table 1**).

Finally, we assessed correlations using Spearman's rank correlation given the ordinal nature of these variables. Task score was not significantly associated with driving frequency ($\rho = 0.156$, $p = 0.311$), or duration of driving experience ($\rho = 0.113$, $p = 0.461$). Likewise, the breadth of gaming experience, defined as the total number of game genres a participant reported playing, was not significantly associated with task scores ($\rho = 0.101$, $p = 0.516$). Furthermore, we examined the correlations between task score and the frequency or duration of play at the individual genre level; no significant correlations were found (all *p*s > 0.2; **Supplementary Table 2**).

**Supplementary Table 1.** Comparison of highway task scores on experience with specific game genres. One person could be included in multiple game genres. The total number of participants was 44. The table shows the mean highway task scores categorized by participants' experience on each game genre. For each genre, an independent-samples Welch's t-test was used to compare the scores between the two groups.

| **Game genre** | **mean score (SD)** | | | |
|---|---|---|---|---|
| | **with experience** | **without experience** | ***t*** | ***p*** |
| Action | 832.59 (213.06) | 866.38 (204.08) | -0.43 | .68 |
| Racing | 842.16 (215.53) | 864.91 (202.863) | -0.31 | .76 |
| Sports | 875.04 (207.34) | 852.58 (206.04) | 0.31 | .76 |
| Role-playing | 883.23 (234.16) | 853.25 (199.61) | 0.32 | .76 |
| Others | 884.78 (217.23) | 824.40 (186.34) | 0.98 | .34 |

**Supplementary Table 2.** Correlations between task score and gaming experience metrics within each genre. This table presents the results of Spearman's rank correlation analyses between task score and frequency of play and duration per session.

| **Game genre** | **frequency** | | **duration per session** | |
|---|---|---|---|---|
| | ***ρ*** | ***p*** | ***ρ*** | ***p*** |
| Action | 0.01 | .92 | 0.05 | .74 |
| Racing | 0.04 | .80 | 0.06 | .71 |
| Sports | 0.07 | .64 | 0.05 | .75 |
| Role-playing | 0.02 | .87 | 0.04 | .77 |
| Others | 0.15 | .32 | 0.19 | .21 |

#### Association between motor ability and task performance

To investigate the association between motor ability and highway task performance, an exploratory analysis was conducted. Data from a total 44 participants were included in the analysis. Motor ability was quantified using five features derived from the fingertapping task: (1) *Initial performance*: the number of correct chunks in the first trial, (2) *Final performance*: the number of correct chunks in the last trial, (3) *Performance improvement*: the relative improvement in performance, calculated as the absolute gain (mean number of correct chunks in last three trials - that of first three trials) normalized by the mean of the first three trials, (4) *Overall accuracy*: the proportion of correct chunks relative to across trials (number of correct chunks / total number of attempted chunks), (5) Total attempts: number of total typed numbers, regardless of accuracy, (6) *Slope*: the slope from linear regression fitted to the number of chunks across 12 trials.

Given the intercorrelation among these features (**Supplementary Fig. 1**), a feature selection process was first performed using Lasso regression with 5-fold cross-validation to mitigate potential multicollinearity. All features were standardized prior to the Lasso analysis. Three predictors identified non-zero coefficients: *Final performance*, *Performance improvement*, and *Overall accuracy*. To examine whether the selected features could predict highway task performance, a multiple linear regression analysis using ordinary least squares (OLS) was conducted using the three predictors. None of the individual predictors significantly predicted task performance (all *p*s > .1). The results indicate the absence of a clear linear association between motor ability and highway task performance (**Supplementary Table 3**).

In addition, we examined whether BIS score predicted highway task performance. A simple linear regression revealed that BIS was not a significant predictor (Model 1; $\beta$ = -13.173, $p$ = .68). To control for the potential influence of motor ability, a multiple regression was conducted including the three fingertapping features as covariates. In this model, BIS remained a non-significant predictor (Model 2; $\beta$ = 0.684, $p$ = .98), indicating that BIS did not predict highway task performance even after accounting for individual differences in motor ability (**Supplementary Table 4**).

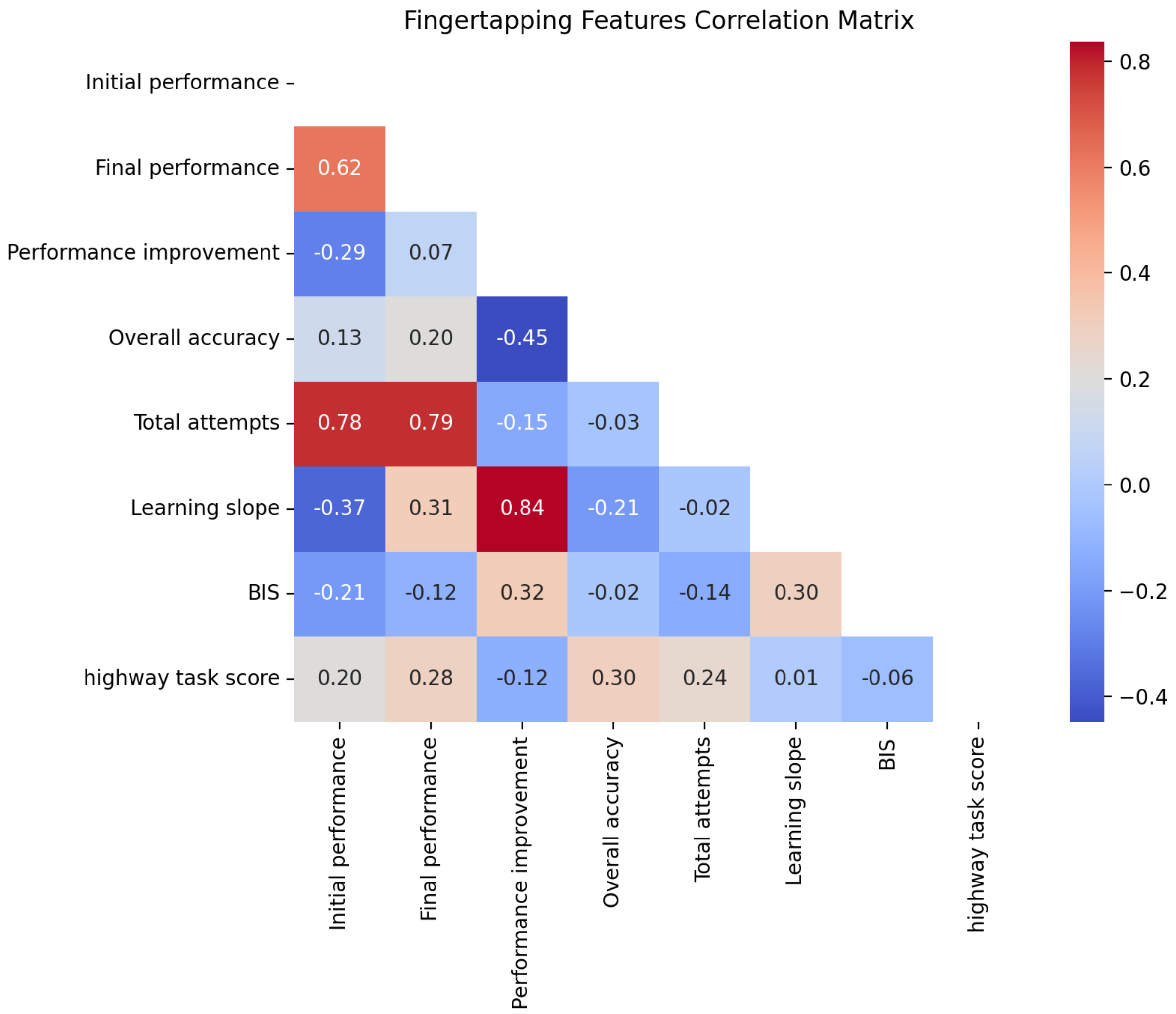


**Supplementary Figure 1.** Correlation matrix of fingertapping variables with BIS and mean highway task score. Correlations were computed with Pearson correlation.

**Supplementary Table 3.** Results of the multiple linear regression model predicting the mean highway task score. Predictors selected from lasso regression were standardized and included into the OLS regression model. $F(3,40) = 2.415$, $p = .081$, adj-$R^2 = 0.090$.

| | $\beta$ | **95% CI** | $t$ | $p$ |
|---|---|---|---|---|
| Overall accuracy | 42.263 | [-24.907, 109.432] | 1.272 | .21 |
| Performance improvement | -24.982 | [-90.880, 40.916] | -0.766 | .45 |
| Final performance | 50.956 | [-11.528, 113.441] | 1.648 | .11 |

**Supplementary Table 4.** Results of the regression models predicting the mean highway task score with BIS and fingertapping task features. Model 1: simple linear regression; Model 2: multiple linear regression including fingertapping task features. Model 1: $F(1,42) = 0.172$, $p = .68$, adj-$R^2 = -0.020$); Model 2: $F(4,39) = 1.767$, $p = .16$, adj-$R^2 = 0.067$.

| | $\beta$ | **95% CI** | ***t*** | ***p*** |
|---|---|---|---|---|
| Model 1 | | | | |
| BIS | -13.173 | [-77.233, 50.887] | -0.415 | .68 |
| Model 2 | | | | |
| BIS | 0.684 | [-64.664, 66.031] | 0.021 | .98 |
| Overall accuracy | 42.164 | [-26.565, 110.893] | 1.241 | .22 |
| Performance improvement | -25.220 | [-95.791, 45.350] | -0.723 | .47 |
| Final performance | 51.068 | [-13.151, 115.287] | 1.608 | .12 |

### Supplementary Figures and Tables

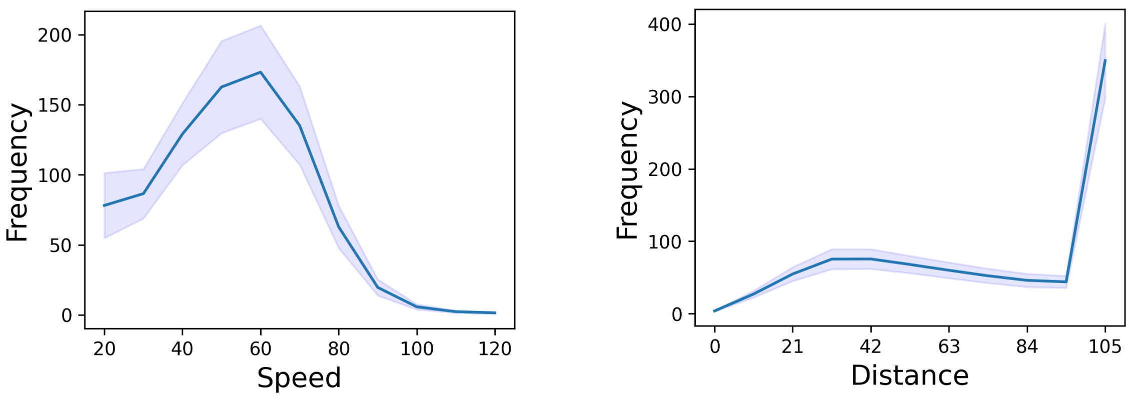


**Supplementary Figure 2.** Frequency distribution of (a) participant's speed and (b) distance from the car ahead in the highway task across all subjects. Speed > 100 accounts for less than 0.4% of the data.

**Supplementary Table 5**. Full list of GLM clusters peaks for overtaking and crash contrasts (N=45). For each event type, subject-level beta maps were submitted to a group-level one-sample t-test against zero. Coordinates (X,Y,Z) are reported in MNI152 space. Voxels were thresholded at $p < 0.01$ Bonferroni-corrected across the whole brain, with a minimum cluster extent of 100 voxels. Sub-peaks that could not be assigned to a labeled region in AAL2, as well as duplicate sub-peaks falling within the same anatomical region as another sub-peak in the cluster, were omitted, retaining the sub-peak with the largest absolute t-value. Cluster peaks that were not labeled in AAL2 were replaced by the sub-peak with the largest absolute t-value within the same cluster (marked with an asterisk).

| | | ***cluster peak*** | | | | |
|---|---|---|---|---|---|---|
| | **ROI** | **X** | **Y** | **Z** | ***t*** | **Cluster Size ($mm^3$)** |
| Overtaking | | | | | | |
| | Cerebellum IX (R) | 10.0 | -56.0 | -44.0 | 15.834948 | 39224 |
| | Superior parietal gyrus (R) | 18.0 | -60.0 | 58.0 | 13.784118 | |
| | Paracentral lobule (L) | -12.0 | -14.0 | 76.0 | 13.779321 | 36768 |
| | Superior frontal gyrus (R) | 26.0 | 2.0 | 60.0 | 12.795658 | |
| | Precentral gyrus (L) | -18.0 | -12.0 | 74.0 | 12.769328 | |
| | Middle frontal gyrus (R) | 30.0 | 0.0 | 52.0 | 12.223913 | |
| | Middle occipital gyrus (L) | -28.0 | -82.0 | 22.0 | 14.733875 | 24712 |
| | Superior parietal gyrus (L) | -30.0 | -58.0 | 58.0 | 13.459487 | |

| | | | | | |
|---|---|---|---|---|---|
| Caudate (L) | -8.0 | 10.0 | 4.0 | 11.614848 | 13560 |
| Putamen (L) | -22.0 | 6.0 | -10.0 | 11.185341 | |
| Inferior frontal triangular (R) * | 36.0 | 22.0 | 10.0 | 9.809741 | 8416 |
| Putamen (R) | 24.0 | 12.0 | 6.0 | 9.724011 | |
| Caudate (R) | 18.0 | 18.0 | 4.0 | 9.383140 | |
| Precuneus (L) | -6.0 | -56.0 | 42.0 | -8.337656 | 2680 |
| Precuneus (R) | 2.0 | -50.0 | 36.0 | -8.137155 | |
| Middle occipital gyrus (R) | 40.0 | -66.0 | 4.0 | 9.673705 | 1400 |
| Cerebellum VIII (L) | -22.0 | -52.0 | -48.0 | 9.742180 | 1248 |
| Angular gyrus (L) | 46.0 | -62.0 | 26.0 | -8.051478 | 872 |
| Parahippocampal gyrus (L) | -28.0 | -32.0 | -18.0 | -9.515575 | 840 |
| Hippocampus (L) | -30.0 | -22.0 | -18.0 | -8.182460 | |
| Insula (R) | 42.0 | -14.0 | 14.0 | -9.695752 | 816 |
| Crash | | | | | |
| Fusiform (R) | 36.0 | -66.0 | -14.0 | 16.976770 | 102544 |
| Hippocampus (R) | 24.0 | -30.0 | -4.0 | 15.006371 | |
| Cuneus (L) | -6.0 | -86.0 | 28.0 | 14.609423 | |
| Calcarine fissure (L) | -6.0 | -74.0 | 16.0 | 13.787267 | |
| Postcentral gyrus (R) | 42.0 | -24.0 | 48.0 | 10.633245 | 11720 |
| Putamen (L) * | -18.0 | 6.0 | -10.0 | -13.253072 | 9800 |
| Caudate (L) | -10.0 | 6.0 | -12.0 | -10.337922 | |
| Putamen (R) | 22.0 | 8.0 | -10.0 | -15.414958 | 8696 |
| Anterior cingulate cortex (L) | -6.0 | 40.0 | 18.0 | 9.621293 | 4808 |
| Middle cingulate cortex (L) | -2.0 | 18.0 | 36.0 | 7.373085 | |
| Superior frontal gyrus (medial; L) | -2.0 | 48.0 | 32.0 | 6.964084 | |
| Superior frontal gyrus (medial; R) | 8.0 | 30.0 | 58.0 | 11.200533 | 3832 |
| Supplementary motor area (R) | 8.0 | 22.0 | 60.0 | 9.127370 | |
| Supplementary motor area (L) | 0.0 | 18.0 | 60.0 | 8.718455 | |
| Middle cingulate cortex (L) | -6.0 | -38.0 | 48.0 | 10.062043 | 3152 |
| Middle cingulate cortex (R) | 4.0 | -32.0 | 42.0 | 8.177912 | |
| Middle cingulate cortex (R) | 6.0 | -20.0 | 30.0 | 10.460085 | 3088 |
| Rolandic Operculum (R) | 40.0 | -12.0 | 20.0 | 9.745925 | 2520 |
| Insula (R) | 36.0 | -16.0 | 12.0 | 9.305426 | |
| Postcentral (L) | -26.0 | -40.0 | 74.0 | 9.840838 | 2360 |
| Supramarginal gyrus (L) | -60.0 | -52.0 | 32.0 | 12.369618 | 2032 |

| | | | | | |
|---|---|---|---|---|---|
| Paracentral lobule (L) | -14.0 | -20.0 | 72.0 | -8.871484 | 1704 |
| Precentral gyrus (L) | -30.0 | -12.0 | 54.0 | -8.126646 | |
| Angular gyrus (R) | 36.0 | -62.0 | 48.0 | 9.398784 | 1592 |
| Insula (L) | -36.0 | -18.0 | 10.0 | 9.051588 | 1000 |
| Rolandic Operculum (L) | -44.0 | -22.0 | 18.0 | 8.073177 | |
| Inferior parietal gyrus (L) | -36.0 | -66.0 | 48.0 | 7.455056 | 808 |

**Supplementary Table 6.** Full list of beta coefficients for the elastic net model predicting full trajectory. Coefficients in the top 10% by absolute value are marked with an asterisk. For Cerebellum Crus I–II, Cerebellum III–X and Vermis I–X, subregions were combined, and the minimum and maximum beta values across subregions are reported.

| ROI | Beta Coefficient (left/right) |
|---|---|
| Precentral gyrus | -0.000633/0.000062 |
| Superior frontal gyrus | 0.000233/0.000179 |
| Middle frontal gyrus * | -0.001081/0.000270 |
| Inferior frontal opercular | left: 0.000482 |
| Inferior frontal triangular | 0.000333/0.000208 |
| IFG pars orbitalis | -0.000051/0.000016 |
| Rolandic Operculum | 0.000036/-0.000321 |
| Supplementary motor area | 0.000011/0.000355 |
| Olfactory cortex | -0.000037/0.000064 |
| Superior frontal gyrus (medial) | 0.000372/-0.000095 |
| Superior frontal gyrus (medial orbital) | -0.000139/0.000289 |
| Gyrus rectus | -0.000090/0.000059 |
| Medial orbital gyrus | 0.000369/-0.000105 |
| Anterior orbital gyrus | -0.000032/0.000014 |
| Posterior orbital gyrus | -0.000058/0.000037 |
| Lateral orbital gyrus | 0.000183/0.000198 |
| Insula | 0.000326/0.000597 |
| Anterior cingulate cortex | -0.000228/0.000044 |
| Middle cingulate cortex | -0.000384/-0.000068 |
| Posterior cingulate cortex | -0.000364/-0.000345 |
| Hippocampus | -0.000274/-0.000770 |
| Parahippocampal gyrus | 0.000093/-0.000121 |
| Amygdala | -0.000092/-0.000153 |
| Calcarine fissure * | 0.000370/-0.001044 |

| | |
|---|---|
| Cuneus * | -0.000748/0.000046 |
| Lingual gyrus | 0.000272/0.000215 |
| Superior occipital gyrus | 0.000599/0.000602 |
| Middle occipital gyrus * | -0.000597/0.001309 |
| Inferior occipital gyrus | -0.000645/-0.000415 |
| Fusiform gyrus * | -0.000012/0.001602 |
| Postcentral gyrus * | -0.000960/0.001146 |
| Superior parietal gyrus | 0.000605/-0.000426 |
| Inferior parietal gyrus | 0.000282/-0.000125 |
| Supramarginal gyrus | 0.000342/0.000281 |
| Angular gyrus | 0.000022/0.000036 |
| Precuneus | 0.000681/0.000249 |
| Paracentral lobule | -0.000611/0.000144 |
| Caudate | 0.000635/0.000364 |
| Putamen * | -0.000316/-0.000838 |
| Pallidum | -0.000077/0.000026 |
| Thalamus | right: -0.000296 |
| Heschl's gyrus | -0.000257/-0.000062 |
| Superior temporal gyrus | -0.000037/-0.000270 |
| Temporal pole (superior temporal gyrus) | 0.000054/-0.000093 |
| Middle temporal gyrus | -0.000399/0.000486 |
| Temporal Pole (middle temporal gyrus) | left: -0.000052 |
| Inferior temporal gyrus * | -0.000143/-0.002128 |
| Cerebellum Crus I–II | left: [-0.000665, 0.000705]<br>right: [0.000038, 0.000161] |
| Cerebellum III–X * | left: [-0.000186, 0.000963]<br>right: [-0.000672, 0.000781] |
| Vermis I–X | [-0.000137, 0.000309] |

**Supplementary Table 7.** Correlations between Neurosynth-based reward BOLD signals and BOLD signals in selected ROIs. The same elastic net pipeline used in the main analysis was repeated with the IRL reward replaced by the mean BOLD signal across voxels within the Neurosynth meta-analytic "reward" mask. Correlations from regions that had non-zero beta coefficients and significant correlation (Bonferroni-corrected $p < 0.01$) are shown. The number of significant ROIs was markedly reduced compared to the main analysis, although the functional associations were consistent. Negative correlations were observed in the orbitofrontal cortex only in crash, and positive correlations were observed in the putamen, instead of the caudate in overtaking.

| ROI | Overtaking (left/right r) | crash (left/right r) |
|---|---|---|
| Olfactory cortex | right: 0.81 | — |
| Amygdala | left: 0.82 | — |
| Inferior occipital gyrus | right: -0.93 | left: 0.85 |
| Putamen | 0.85/0.85 | — |
| Pallidum | left: 0.85 | — |
| Frontal medial orbital cortex | — | right: -0.80 |
| Gyrus rectus | — | left: -0.90 |
| Medial orbitofrontal cortex | — | right: -0.91 |
| Anterior orbitofrontal cortex | — | right: -0.87 |
| Lateral orbitofrontal cortex | — | left: -0.80 |
| Supramarginal gyrus | — | right: 0.77 |
| Paracentral lobule | — | right: 0.79 |
